# Pleistocene dispersion supports a unique native diversity in the Colombian avocado germplasm

**DOI:** 10.1101/2023.01.27.525883

**Authors:** J.A. Berdugo-Cely, A.J. Cortés, F. López-Hernández, P. Delgadillo, I. Cerón-Souza, P. Reyes-Herrera, A. Navas, R Yockteng

**Author notes:** **Correspondence**, Cortés, A.J and Yockteng, R., AGROSAVIA, Colombia. and.

## Abstract

Genomic characterization of ex-situ collections optimizes the utilization of genetic resources, identifies redundancies among accessions, captures cryptic variation, establishes reference collections, and ultimately assists pre-breeding and breeding efforts. However, the integration of population genomic analyses is often lacking when studying the biodiversity of crop gene pools. Here, we present modern classification and machine learning approaches to characterize and harness the genebank of an agrobiodiversity hotspot on *Persea americana* Mill., an iconic American fruit tree crop that has seen an unprecedented expansion worldwide. We selected 144 accessions from the Colombian National genebank and 240 materials from local plantations in the Colombian Northwest Andes. We genotyped them using a strategy based on reduced representation sequencing. We included available sequences of genotypes from known avocado races, Mexican, Guatemalan, and West Indian, to discover SNPs, analyze the population structure and identify possible new genetic groups in Colombian germplasm. We detected a population structure suggestive of a new fourth race in Colombia, with a possible genetic substructure related to geographical origin (Andean and Caribbean). Hybrid determination and ABC modeling suggested rampant inter-race geneflow. They supported the hypothesis of the high mobility of native avocado trees from Central America to Northern South America starting in the Pleistocene. Our study supports that Colombia might be a new diversity center for *P. americana.* Genotypes of the two newly identified Colombian groups can be used as parents in plant breeding strategies to generate cultivars adapted to specific ecogeographical regions of Colombia.

## Introduction

Studying plant variation in crop wild agrobiodiversity hotspots is a significant research avenue for discovering novel attributes that may increase worldwide food and nutrient requirements (Ramirez-Villegas et al. 2022). Avocado (*Persea americana* Mill., Lauraceae) is one of the priority crops in highly diverse Mesoamerican and South American regions, dating to use and selection at least 9,000 years near of its center of origin in Mexico and Central America (Smith 1966, 1969). During pre-Columbian times, the obligate route for exchanging crops and wild relatives between Mesoamerica and South America was through northwest South America (Larranaga et al. 2021) Thus, when the Spanish arrived in America, avocado distribution ranged from Mesoamerica to Ecuador and northern Peru (Wolters 1999; Galindo-Tovar et al. 2007, 2008). Therefore, Mesoamerica and the north of South America are considered a diversity hotspot for tree species from the Lauraceae family (Pironon et al. 2020), but its genetic diversity remains unexplored (Chen et al. 2009; Galindo-Tovar and Arzate-Fernández 2010). In Colombia, native avocado production is mainly intended for the internal consumption of fresh and processed fruit. Despite this, the northern Colombian Andes present favorable agro-climatic conditions (from 0 to 2,200 m.a.s.l.) to produce avocado to be exported at the end of the year when the worldwide demand is unsatisfied (Rios Castaño and Tafur Reyes 2003). Recent exports to Europe, Japan, and the USA, among other countries, totalized 146 million USD in 2020, reinforcing Colombia’s prospect of supplying avocado markets abroad (Statistica 2022). In 2020, Colombia reached a second place as an avocado producer with 876.7 thousand tons of global production (FAOSTAT 2022), after Mexico, which remains the first producer and consumer (Galindo-Tovar et al. 2011; FAOSTAT 2022).

Avocado originated in Central America and Mexico (Arumuganathan and Earle 1991; Alcaraz and Hormaza 2007; Calderón-Vázquez et al. 2013), probably in the Tehuacan Valley in the state of Puebla (Galindo-Tovar et al. 2007). The genus *Persea* Mill. has two different subgenera, *Persea* and *Eriodaphne,* each containing 12 and 11 species, respectively (van der Werff 2002). The species of *Persea* subgenus, to which avocado (*P. americana*) belongs, are distributed in Central America and Northern South America. Avocado’s distribution extended until Peru and continued to be mediated by human populations following commercial pre-Columbine routes (Bergh and Ellstrand 1986; Wolters 1999; Galindo-Tovar et al. 2008). The *Eriodaphne* subgenus (so-called aguacatillos) contains species resistant to root rock disease caused by *Phytophthora cinnamomi,* the most important disease of avocado. However, gametes from the *Eriodaphne* subgenus and *P. americana* are not compatible (Zentmyer and Schieber 1992).

The Avocado species currently comprises three horticultural races that differ in origin, genetic diversity, and commercial characteristics, Guatemalan [*P. americana* var. guatemalensis (L.) Wms.], Mexican [*P. americana* var. drymifolia (Schlecht. et Cham.) Blake] and West Indian [*P. americana* var. americana Mill.)] (Rendón-Anaya et al. 2019). The Guatemalan race originated in the mid-altitude highlands of Guatemala and is characterized by having tiny seeds and late fruit maturity. The Mexican race originated in the mid-altitude highlands of Mexico but exhibited early fruit maturity and cold tolerance. In contrast, the West Indian race originated in southern Mexico and Central America’s lowlands and has large fruits with low oil content (Rendón-Anaya et al. 2019). However, this race has open questions about its origin and dispersion routes because Spanish chroniclers described the presence of avocado trees with West-Indies characteristics in northern South America, including Colombia, Ecuador, and Peru at both the Pacific Coast and in the Amazon region (Galindo-Tovar and Arzate-Fernández 2010).

Several genetic studies of avocados supported the three major recognized races and their evolutionary origin. Expressed Sequence Tag (EST), Single Sequence Repeats (SSR), and Single-Nucleotide Polymorphism (SNP), among others (Gross-German and Viruel 2013; Ge et al. 2019; Talavera et al. 2019) were used in these studies. Moreover, several avocado germplasm banks have characterized their accessions at the genetic level to identify the ancestry of the conserved genotypes. They are the Venezuelan gene bank - INIA-CENIAP (Ferrer-Pereira et al. 2017), the National Gene Bank repository (SHRS ARS USDA) in Miami (Boza et al. 2018), and the Spanish Gene Bank (CAÑAS-GUTIÉRREZ et al., 2019).

In 2019, Rendón-Anaya et al. (2019) released the first genome sequence of an avocado Hass cultivar providing a reference of 980 Mb in length. The genomic information reinforced race substructure and provided evidence for the hybrid origin of commercially essential varieties such as Mexican/Guatemalan avocado var. Hass (Rendón-Anaya et al. 2019). Although avocados growing genomic resources promise to aid the conservation, breeding, and discovery of new commercial varieties, avocado germplasm from South American Andean tropics such as Colombia is still poorly characterized, and accessions and seedling rootstocks lack race classification. Therefore, the goal of this study was to combine the reported SNP diversity of avocado races with the genotyping of commercials and native accessions from the Colombian avocado germplasm bank and different seedling rootstocks from commercial plantations via a high throughput reduced representation library sequencing method.

Specifically, we focused on the following questions: (1) does the Colombian avocado germplasm belong to the recognized races, or could they represent a new biodiversity center for *P. americana?* (2) which demographic scenario better explains the genetic structure of Colombian germplasm according to possible dispersal scenarios of avocado from Mesoamerica to Colombia? (3) to which races or genetic groups belong the rootstocks nowadays found in commercial plantations in the Colombian North Andes? Exploring the questions above would inform whether Colombian avocado resources may provide novel allelic diversity beyond the classical races in Central America.

## Material and methods

### Plant Material and sequences of genomic resources

This study analyzed the genetic diversity of 456 avocado samples (*P. americana)* from two sources: the Colombian Germplasm Bank - CGB (n=144) and AGROSAVIA’s pre-breeding elite collection – EC (n=240) (Table S1). Additionally, sequences of genomic resources of seventy-one (71) individuals from the study of Talavera et al. (2019) representing the three recognized avocado races (Guatemalan-GU; Mexican-ME, and West Indian-WI) and hybrids among the races were included as Reference Materials (RM). At the same time, the genome sequence of *P. schiedana* was used as an Outgroup (O) (Rendón-Anaya et al. 2019).

The CGB is conserved at the Colombian Corporation of Agricultural Research (AGROSAVIA) at the Palmira research station in the province of Valle del Cauca (3°32’22.0”N76°18’13.0”W, 1,000 masl). Of the 144 accessions, fifteen are commercial varieties or cultivars, and 129 correspond to Colombian native and criollo trees (native x introduced hybrids). Some accessions were sampled in 2006 from contrasting Colombian agroclimatic regions, particularly in high-soil humid regions, where it is probable to find tolerant materials to *P. cinnamomi* (Rodriguez Henao 2015). Following the Bioversity manual for avocados, this germplasm has been characterized by juvenile morphological and botanical traits (IPGRI 1995). The EC has 240 seedling rootstocks from eight commercial plantations in the Antioquia Department, Colombia’s primary avocado-producing region (Table S1). This germplasm has a previous genetic description using SSR markers (Reyes-Herrera et al. 2020).

### Molecular genetic characterization of CGB and EC materials

#### DNA Isolation

We extracted genomic DNA from foliar tissues of 144 accessions conserved in the CGB and from root tissues of 240 seedling rootstocks (EC) using the DNeasy Plant Mini Kit (QIAGEN, Germany) with the following modifications: first, we added 450 μL of the AP1 solution to an equal volume of 20% SDS (Sodium Dodecyl Sulfate), and we incubated the samples for 30 mins at 65 °C and froze for 60 mins at −20 °C. DNA quality was verified by electrophoresis in 1% agarose gels dyed with Syber Safe (Invitrogen), and the DNA concentrations were measured in a Qubit® 2.0 fluorometer (Life Technologies).

#### Genomic Libraries Construction and Sequencing

Each DNA sample was double digested using *PstI* (CTGCA - 37 °C) and *ApeKI* (G/CWGC - 75 °C) enzymes, following the recommendations of New England BioLabs. Genomic libraries were constructed from the digested DNA using the NEBNext Ultra DNA kit for Illumina (E7103L). To identify the sequences corresponding to each sample, we used the indexes of the 96 NEBNext Multiplex Oligos kit for Illumina (E6609L). Each genomic library was quantified through fluorescence in a Qubit 2.0, and the average size of the fragments of some random samples was determined by digital electrophoresis on a TapeStation 4200 (Agilent). Finally, the genomic libraries were diluted to a concentration of 10nM and pooled in groups of 96 samples. The pooled samples were sequenced using a *paired-end* strategy with 150 sequencing cycles in an Illumina HiSeq X equipment (Macrogen, Inc. Korea). The demultiplexing of sequences for each genotype was implemented using the bcl2fastq Illumina software (bcl2fastq 2022). Raw data sequences were deposited in the Sequence Read Archive (SRA) on the National Center for Biotechnology Information (NCBI) database in the Bioproject number PRJNA878519.

#### SNPs identification among the CGB, EC, RM, and O samples

To discover SNPs among all samples, we made the SNP calling using the pipeline reported by Osorio-Guarín et al. (2020). In summary, for all sequence files of each analyzed sample (downloaded and generated in this study), we used the FastQC software (Andrews 2010) to verify the quality of the Fastq files of each sample. The adapter’s primers and sequences with quality less than Q30 and length less than 50 base pairs (bp) were filtered using Trim-Galore software (Krueger 2012). The Burrows-Wheeler Aligner (BWA) software (Li and Durbin 2009) was used to align the sequences against the *P. americana* Hass cultivar reference genome version 2.0 (~913 Megabases - Mb) (Rendón-Anaya et al. 2019). We used Picard (Picard 2022) and the Genome Analysis ToolKit - GATK (GATK 2022) software to remove PCR duplicates, to correct mapping quality assignment, and to realign and recalibrate the reads. The SNP calling was done using the *Unifiedgenotyper* option of the GATK. Finally, we used the VCFtools software (Danecek et al. 2011) for filtering the SNP markers with a Minimum Allele Frequency (MAF) under 5%, a minimum of 2X of Sequencing Depth (SD) per position, maintaining biallelic markers and excluding InDels (Insertion-deletion) markers. Samples and SNP markers with percentages higher than 20% (genotype level) and 5% (marker level) of missing data were excluded.

The nucleotide sequences were obtained from different tissues and genomic resources (Rendón-Anaya et al., 2019; Talavera et al., 2019). Then, the final filter for polymorphic SNP markers discovery was implemented to get four different datasets used in further analyses (Table 1). Metrics related to SNP markers such as Polymorphic Information Content (PIC), Minor Allele Frequency (MAF), Nucleotide Diversity (ND), Mean Depth of Sequencing (MDS), Genetic Diversity (GD), Observed Heterozygosity (Ho), and Transition/Transversion (Ts/Tv) statistics were determined in the dataset 3 using VCFtools software (Danecek et al. 2011) and *SNPready* (Granato et al. 2018) in R package (R 2022).

**Table 1.** SNP markers in Avocado datasets generated to implement the analyses proposed in this study. **CGB**: Colombian Germplasm Bank; **EC:** Elite collection; **RM:** Reference Materials; **O:** Outgroup.

| Number of datasets | Dataset included | Number of samples post filters | Number of polymorphic markers | Implemented Analysis |
| --- | --- | --- | --- | --- |
| 1 | CGB + RM | 199 | 4931 | Population Structure/Demographic scenarios |
| 2 | CGB + RM + O | 203 | 3899 | Phylogenetics |
| 3 | CGB + Pure RM | 154 | 26048 | Ancestry in CGB |
| 4 | EC + Pure RM | 289 | 227 | Ancestry in EC |

#### Genetic Statistical Analysis

To answer the research questions proposed in this study about the classification of the Avocado Colombian germplasm in the reported races (ME, GU, and WI) and the presence of new genetic groups, we used four different approaches as described below.

#### Unsupervised non-parametric Genetic Cluster Analysis

Population stratification was characterized using dataset 1, which included 4931 SNPs identified in 199 samples (Table 1) using two complementary non-parametric approaches related to unsupervised machine learning: partitioning and hierarchical clustering. We had to perform a prior step to reduce the dimensionality of the SNP dataset through Principal Component Analysis (PCA) (Alhusain and Hafez 2018; Foote et al. 2019) using *R-glPca* function in the *R-adegenet* package (Jombart and Ahmed 2011). Once dimensionality was reduced, we applied unsupervised non-parametric clustering approaches. For partitioning clustering, we used K-means (MacQueen 1967; Lloyd 1982), Partitioning Around Medoids (PAM), and Clustering Large Applications (CLARA) (Kaufman and Rousseeuw 2009) algorithms, and for hierarchical clustering, we ran AGglomerative NESting (AGNES) y DIANA algorithms (Divisive analysis) (Kaufman and Rousseeuw 2009) using the *ward.D2* method in R software (R 2022).

To find the optimal number of subpopulations or K Clusters for each unsupervised non-parametric clustering algorithm, we conducted in R the two complementary methods *NbClust* (Charrad et al. 2014) and *optCluster* (Sekula et al. 2017). First, we ran the *NbClust* algorithm, an internal measure’s function that integrates 30 indexes to determine the optimal K value based on *Ward.D2* method and the Euclidean distance evaluating three (K=3) to eight (K=8) putative subpopulations. Second, we ran the *optCluster* function, an improvement of the traditional *clValid* algorithm (Brock et al. 2008) that optimizes both the K score and the clustering approach based on internal, stability, and biological measures. In this sense, the *optCluster* function validated at once all clustering algorithms by each approach using cross-entropy and algorithmic genetic methods, using weighted Spearman footrule distance (Kumar and Vassilvitskii 2010), evaluating three (K=3) to eight (K=8) putative subpopulations.

#### Statistical Genetic Structure Analysis

To identify the genetic differentiation among detected clusters in the previous section, we conducted an analysis of molecular variance (AMOVA) using 1000 permutations in the *R-poppr* package (Kamvar et al. 2014). Pairwise F_ST_ values were calculated in the *R-dartR* package (Gruber et al. 2018), while the genetic diversity measured by observed (Ho) and expected (He) heterozygosity were calculated in the *R-hierfstat* package (Goudet et al. 2022). Mantel correlations test at 1000 permutations between genetic differentiation and geographic distance within and between pairs of clusters using the *R-geosphere* package (Hijmans et al. 2021). Diagrams were drawn in the same software depicting pairwise F_ST_/(1–F_ST_) vs. geographic distance (Km), following Rousset, (1997). Meanwhile, bidirectional gene flow among pairs of populations was estimated as the number of migrants per generation (N_e_m) following the methodology described by Beerli & Felsenstein, (1999), which is the inverse surrogate of pairwise F_ST_ scores. We draw, using the *R-qgraph* package (Epskamp et al., 2012), the networks depicting pairwise bidirectional migration rates across all datasets (Epskamp et al. 2012).

#### Reconstruction of demographic scenarios

We used the DIYABC Random Forest v1.0 program (Collin et al. 2021), an extended version of DIYABC v2.1.0. This tree-based classification method uses supervised machine learning to classify demographic evolutionary scenarios and estimate their parameter robustness based on a certain number of simulations per scenario using the genetic groups obtained in the previous analysis. We simulated 120,000 sets for the 4931 selected loci of dataset 1 (Table 1), and 1000 trees for model choice and 1000 as the number of out-of-band testing samples to estimate the historical parameters given the highest posterior probability model from the eight simulated. Because there is no experimental estimate of the SNP mutation rate available for avocados, the mean mutation rate followed an *a priori* uniform distribution. Each locus had a possible range of two allelic states, as expected for bi-allelic SNP markers in a diploid species. Moreover, we set the divergence in the number of generations as equal or higher than in this order: *t*_3_ ≥ *t*_2_ ≥ *t*_1_. First, we intended five groups: the races Mexican (ME), Guatemalan (GU), West Indian (WI), and the two Colombian groups identified. However, several runs testing these five groups and settled parameters never showed convergence (data not included). Therefore, using previous parameters, we simplified the analysis into four groups: the three races (i.e., ME, GU, and WI) and a single Colombian group with accessions from the two subgroups named CO, which showed convergence.

The eight scenarios tested were following. Scenario 1 (Fig. S1a) was the null hypothesis, indicating that all the four genetic groups diverged simultaneously at *t*_1_. Scenarios 2-4 considered three divergence times among the four groups (i.e., *t*_1_, *t*_2_, and *t*_3_) but tested what was the most ancient group, considering the two extremes of the distribution, from Mesoamerica to the North of South America. In scenario 2 (Fig. S1b), the most ancient divergence occurred at *t*_3_ and derived CO. Then, at *t*_2_ derived WI, and finally, at *t*_1_, the most recent common ancestor between ME and GU diverged. In scenario 3 (Fig. S1b), CO is derived from the most ancient divergence *t*_3_, GU evolved at time *t*_2_, and at the most recent divergence time, *t*_1_ evolved ME and WI from the most recent common ancestor. In scenario 4 (Fig. S1d), the ME group is the most ancient group derived at *t*_3_, GU derived at time *t*_2_, and at *t*_1_ diverged CO and WI from the most recent common ancestor. Scenario 5 (Fig. S1e) contemplated a potential hybrid evolution. Here, CO is the most ancient group derived at *t*_3_; then, at *t*_2_, WI and ME split; at *t*_1_, GU is derived from the hybridization between WI and ME. Alternatively, scenarios 6-8 simulated the evolution of the most recent common ancestor at *t*_3_ from the four groups, then, the two clades that split at times *t*_2_ and *t*_1_ generating two sister groups. In scenario 6 (Fig. S1f), the sister groups were WI-ME, which split at *t*_2_, and CO-GU that split at *t*_1_. In scenario 7 (Fig. S1g), the sister groups were GU-ME, which split at *t*_2_, and CO-WI, that split at *t*_1_. Finally, in scenario 8 (Fig. S1h), the sister groups were WI-GU, which split at *t*_2_, and CO-ME that split at *t*_1_.

After selecting the best model among the eight tested, we converted the outputs (i.e., means, standard deviation, and confidence intervals at 0.05 and 0.95) about divergence times *t*_1_, *t*_2_, and *t*_3_ expressed in the number of generations (g) (i.e., the average time in years between two consecutive generations, such as parent and offspring) to compare them with two hypothesis about the evolution and dispersion of avocado from Mesoamerica to the Northern South America. The avocado species is a perennial woody tree (Miller and Gross 2011) with a variable generation time depending on plantations and crosses among cultivars. Therefore, the conversion from g to time is biased by the number of generations we settled for avocado. To acknowledge this variability on g times among different studies, we decided to use three different years. The short one g = 10 represents the earliest age of fruiting observed among 11 pairwise cultivar crosses (i.e., ranging between three and 14 years) (Thorp et al. 2015). Also, we used g = 50 and g = 100 because there are plantations with 30 years old trees used in experiments, and reports of ancient avocado trees still alive that produced fruits (Krome 1956; Goodall et al. 1970; Zuazo et al. 2021).

We compared the times when occurred the three splits from the best model with two hypotheses about the evolution and genetic diversification of avocado based on archeological data and Spanish chroniclers’ descriptions (Galindo-Tovar et al. 2007; Galindo-Tovar and Arzate-Fernández 2010). The species of *Persea* subgenus, to which avocado (*P. americana*) belongs probably diversified as product of available habitats in the Pleistocene (i.e., from 2’580,000 to 11,700 years ago) (Galindo-Tovar et al. 2007). If it so, dispersion and diversification occurred mediated by big mammals and hunter-gatherers’ humans as they migrated from Mesoamerica to the Northern South America through the Central American Isthmus (Galindo-Tovar et al. 2007). In contrast, if the dispersal and diversification of avocado occurred during agriculture and village life, we expected that the splits occurred during the Holocene, the most recent period of the Quaternary that began approximately 11,650 years ago until today (Krome 1956; Goodall et al. 1970; Galindo-Tovar et al. 2007; Miller and Gross 2011; Thorp et al. 2015; Zuazo et al. 2021).

#### Maximum Likelihood method for Phylogenetic Analyses

The phylogenetic analysis was conducted using dataset 2, which includes 3899 SNP markers for 202 avocado samples (Table 1) and an Outgroup (O), the close relative species *P. schideana.* Although this species could potentially hybridize with avocados (Ashworth and Clegg 2003), it behaved as an outgroup in our analyses. In a preliminary analysis (data not shown) in which we also used the species *Phoebe bournei* as an outgroup, *P. schiedana* was more related to this species than to avocados. Therefore, we decided to use only *P. schiedana* as an outgroup because we could recover more SNPs. The alignment of nucleotide sequences of all targeting SNPs was visualized and verified in Geneious v8.1.7 (Kearse et al. 2012). The phylogenetic tree was reconstructed using the Maximum Likelihood (ML) method in the PhyML v3.1 software (Guindon et al. 2010) with 1000 bootstrap replicates. The Colombian genotypes with geo-referenced information were used to construct a geo-referential map using the *raster* (Hijmans 2022), and *ggplot2* (Wickham 2022) R-packages.

#### Ancestry Analyses

To determine which races of avocado are present in the CGB and EC, we ran two independent ancestry analyses using datasets 3 and 4 (Table 1) because each matrix presented a different number of polymorphic markers, 26048 and 227, respectively (Table 1). These datasets included genotypes with high genetic ancestry (>80%) to one genetic race (ME, GU, WI) and the Colombian group (CO) determined in the Unsupervised non-parametric Genetic Cluster Analysis. In the case of the CO group, we selected samples representing the two possible subpopulations detected in the Colombian collection (CO_Andean and CO_Caribbean) through phylogenetic analyses. The samples with a clear genetic ancestry for these five genetic clusters were used as reference genotypes to implement a supervised analysis in *ADMIXTURE* software (Alexander and Lange 2011). These analyses were run for a K of five (ME, GU, WI, CO_Andean, and CO_Caribbean) using a Cross-Validation (CV) of 10. CGB and EC samples with >80% of ancestry were assigned to a specific genetic cluster, and mixed samples (< 79%) were cataloged based on their admixture proportion (*e.g.,* CO_Andean x CO_Caribbean).

## Results

### Number and statistics of SNP discovered in Avocado datasets

The sequencing of the 384 avocado genotypes of CGB and EC generated a total of 3.132.393.704 sequence reads (Table S1) that were merged with the available sequences for the 71 reference genotypes (Talavera et al. 2019), and the outgroup, *P. schideana* (Rendón-Anaya et al. 2019). Four SNP datasets were generated, and each included different samples (CGB, EC, RM, and O) depending on the analyses. The SNPs ranged from 227 in dataset 4, which included EC and Pure RM genotypes, to 26048 in dataset 3, which contained CGB and Pure RM genotypes (Table 1). Considerable differences in the number of SNPs were observed in datasets, including EC genotypes, from which the DNA was obtained from root tissue. However, the number of sequences generated in the CGB (3 million – M to 52M) and EC (1.3M to 56M) genotypes was similar (Table S1). Then, a high number of sequences from avocado root tissue was generated, but the percentage of missing data among the samples was high in the SNP calling. It has been reported that the efficiency of the nucleic acid extraction process differs from the tissue used in DNA extraction processes because the physical (lignified grade) and chemical composition (higher presence of metabolite secondaries) and diameter of the root tissue can affect these processes. Therefore, when working with root tissue in genomic approaches, it is recommended to use its tissue in the same conditions (dryness) and characteristics (size) (Fisk et al. 2010; Zeng et al. 2015; Oliveira et al. 2015; Miller et al. 2017). However, the number of SNPs identified in this study demonstrates *a priori* the degree of differentiation and genetic diversity present among the evaluated germplasm, which is correlated with the average values of the PIC (0.22), Ho (0.27), and MAF (0.19), being theses higher than the results reported by Talavera et al. (2019).

The statistics of SNPs were calculated using dataset 3 (Table 1), which had the highest SNP number (26048) mapping in 8082 contigs of the Hass genome reference. This dataset had a Ts/TV proportion of 1.27, with 15487 transitions (Ts) and 10561 transversions (Tv) substitutions and an MDS mean of 5.25 (SD=10.49) (Fig. S2a). The polymorphisms had a mean MAF of 0.19 (SD=0.13), ranging from 0.05 to 0.5 (Fig. S2b), and most were highly informative, with a mean PIC of 0.22 (SD=0.096), ranging between 0.1 to 0.4 (Fig. S2c). The GD and ND statistics presented similar values to means (0.27; SD = 0.13) and ranges (0.01-0.5), and the major proportion of SNPs showed ranges between 0.1 to 0.2 (Fig. S2d and S2e). Ho values ranged between 0 to 1.1 (Fig. S2f).

### Avocado Genetic Stratification was Consistent with Four Subpopulations

The two unsupervised non-parametric clustering approaches systematically exhibited four clusters as subpopulations (Table S2 and S3). Initially, we obtained three principal components: the first component explained the 26.77%, the second component explained the 12.98%, and the third component explained the 6.93% variance from raw SNPs. Based on the principal component, the *NbClust* function provided four optimal clusters by running the *ward. D2* method (Table S2). Meanwhile, independent of the clustering approaches or validation method as cross-entropy and genetic algorithms, the *optCluster* function recurrently suggested four optimal clusters. PAM was the best partitional clustering method (*Cross-Entropy score:* 25.81552, *Genetic Algorithm score:* 25.49845), and AGNES was the best hierarchical method (*Cross-Entropy score*: 20.81848, *Genetic Algorithm score:* 20.81848). In this way, the *optCluster* function suggested PAM as the best algorithm compared to AGNES. Thus, we assigned each accession to an assigned cluster that was distributed in four groups: GU Cluster (n=43), CO Cluster (n=79), ME Cluster (n=19), and WI Cluster (n=58) (Fig. 1). This analysis showed the separation of CO Cluster in the first component. The Mexican-ME, Guatemalan-GU, and West Indian-WI races from Talavera et al. (2019) composed the other three clusters. The Colombian group-CO was exclusively composed of Colombian native accessions. The other three groups included the RM, the admixtures between Mesoamerican genotypes (*e.g.,* GUxWI, GxM, MxWI), Commercial Materials (CM), and CGB accessions with high ancestry of Mesoamerican races (ME, GU, and WI) (Fig. 1).

**Fig. 1** Principal Component Analysis (PCA) of 199 avocado accessions (CGB and RM) with 4931 SNPs. The colors indicate the assignment to a genetic group based on the Partitional Around Medoids (K-medoids). The black letters indicate the three recognized avocado races (GU: Guatemalan, ME: Mexican, and WI: West Indian). Commercial Materials (CM).

### Genetic Divergence was Concordant with Botanical or Spatial Differentiation

A molecular variance analysis (AMOVA) indicated that 54.9% of the genetic variation was between the four clusters determined in the previous section, while 45.1% was found within the clusters. This result indicated a solid genetic structure supported by a global Phi value of 0.548 (*p*=0.001). Pairwise F_ST_ scores ranged from 0.207 (ME Cluster *vs.* GU Cluster) to 0.489 (ME Cluster *vs.* CO Cluster) supporting a robust genetic differentiation between the Colombian group (CO) and the Mesoamerican races with pairwise F_ST_ of 0.489 (ME Cluster *vs.* CO Cluster), 0.466 (GU Cluster *vs.* CO Cluster), and 0.297 (WI Cluster *vs.* CO Cluster) (Fig. 2a). The observed (Ho) and expected (He) heterozygosity showed that Mesoamerican groups are in general more heterozygous, (WI Ho=0.277 and He=0.244), ME (Ho=0.241 and He=0.208), and GU (Ho=0.215 and He=0.194), than the CO group (Ho=0.178 and He=0.166). The estimates of bidirectional migration rates (N_e_m) among subpopulations were low (≤ 1) and ranged between 0.29 and 0.95. Estimated migration rates were asymmetric among the four genetic clusters and, as expected from the F_ST_ comparisons, were lower (0.38 ± 0.11 *vs.* 0.77 ± 0.11) in the pairwise contrast with the subpopulation from the Colombian group (ranging 0.26 - 0.59) (Fig. 2b). We further explored the genomic landscape of divergence in avocados as a function of geographic (IBD) differentiation. Explicit correlations of genetic distance with geographic gradients, obtained using Mantel tests within and between pairs of clusters, were not significant given 1000 permutations (Fig.S3a-S3d).

**Fig. 2** Pairwise F_ST_ **(a)** and bidirectional migration rates (N_e_m) **(b)** values among the four Avocado genetic clusters determined by population genetic structure analysis. In the Figure, GU Cluster corresponds to the Guatemalan genetic group, CO Cluster corresponds to the Colombian group, ME Cluster genetic corresponds to the Mexican genetic group, and WI Cluster corresponds to the West Indian genetic group.

### Ancient Colonization of the Northern Andes Predated Recent Diverge in Mesoamerica

We used the DIYABC random forest method to test the best demographic scenario explaining the evolution of the Colombian group (CO Cluster) against the other races (i.e., Mexican-ME, Guatemalan-GU, and West-Indian-WI) (Fig. S1). Scenario 7 best fitted our data with 592 votes out of 1000 random forest trees and a posterior probability of 0.772 (Fig. 3a). This scenario corresponds with the geographic origin of the three races from Mesoamerica and the Colombian group, from the Northern South America. However, it also reveals a novelty, the parallel evolution of two clades of avocado according with their proximity based on geographic location and habitat. One clade represents the recent common ancestor between ME that originated in the mid-altitude highlands of Mexico and GU that originated in the mid-altitude highlands of Guatemala. The other clade, represent the recent common ancestor between WI originated in Southern Mexico and Central American’s lowland and CO from the North of South America, subdivided geographically in an Andean group and in a Caribbean group within Colombia. The number of votes for the other seven scenarios discarded in decreasing order were: Scenario 5 (132 votes), scenario 3 (112 votes), scenario 4 (108 votes), scenario 8 (26 votes), scenario 1 (17 votes), scenario 2 (13 votes) and scenario 4 (zero votes).

Based on scenario 7, we found that all evolutionary divergences using the tree different generation time (g) tested, occurred during the Pleistocene (i.e., between 12000-2’000.000 years ago), supporting that divergence is ancient and occurred before the Holocene. It means before, agriculture and village life started in the Americas. Thus, the older divergence between two ancestral avocado clades (t_3_) occurred between 59,152 years ago, 95% CI: 27,985-90,630 (at g=10) and 591,524 years ago, 95%: 279,803-906,300 (at g=100). Moreover, the divergence between ME and GU races (t2) occurred between 29,256 years ago, 95% CI:11,399-53,860 (at g=10) and 292,560 years ago, 95% CI: 113,998-538,600 (at g=100). In parallel, the divergence between the CO and WI race (t_1_) occurred between 37,608 years ago, 95% CI: 15,680-63,439 (at g=10) and 376,083 years ago, 95% CI: 156,800-634,391 (at g=100) (Fig. 3b).

**Fig. 3** Demographic evolution of avocado reconstructed by the DIYABC random forest methodology. **(a)** The most probable evolutionary scenario of avocados in Mesoamerica and the North of South America showed a posterior probability of 0.772. This scenario suggests a parallel evolution of two ancient clades that diverged at t3. One clade split at t2, originating the Mexican (ME) and Guatemalan (GU) races. The other clade split simultaneously at t1, forming the Colombian group (CO) and the West Indian (WI) race. The divergence times are not to scale. **(b)** The divergence time in years of the best scenario 7, assuming ten years (green), 50 years (orange), and 100 years (violet) of generation times (g). The white circles represent the mean estimates, the thick hash marks indicate the standard deviation, and the thin hash marks indicate the confidence interval at 95%. The grey vertical dot line separates the Pleistocene and the Holocene within the Quaternary era. The Pleistocene started 2’000.000 years ago and ended 11,700 years ago. At that time, the Holocene started until the present.

### Phylogenetic Analysis Supported Stratification Across Four Major Subpopulations

Using the SNP alignment for 202 samples, the phylogenetic tree regrouped most samples in four supported clades, corresponding to the reference groups and a new clade grouping most Colombian genotypes from the AGROSAVIA germplasm bank (Fig. S4). However, some accessions did not fall within a single clade. Based on the clustering analysis, they corresponded to hybrids or admixtures. Thus, we created a new phylogenetic analysis excluding those hybrids or admixtures, with 92 samples and 3899 SNPs (Fig. 4). The analysis confirmed four clades with bootstrap support (BS) of 100% that match the clustering analyses (Fig. 1). The basal clade corresponded to samples belonging to the Mexican (ME) race. The accessions from the Guatemalan (GU) race formed the next clade in position, followed by the clade with the West Indian (WI) materials. The Colombian (CO) germplasm bank accessions and the West Indian (WI) race are related in two supported clades in the most derived phylogeny position. The Colombian (CO) clade is divided into two subgroups. Based on passport data, the avocado genotypes mainly from the Andean region (CO_Andean: Cauca, Huila, Quindio, Risaralda, Santander, Meta, and Tolima departments) were regrouped in a well-supported clade (BS=100%), and in a second weak-supported clade (BS=43%) regrouped samples from the Caribbean region (CO_Caribbean: Bolivar, Cesar, and Magdalena departments). Only six accessions from the Colombian germplasm fell in the Mesoamerican clades, four samples (WALDIN, QUIBU_11, VADU_10, and VADU_14) in the West Indian clade, one (HASS) in the Guatemalan clade, and one (TOPA_TOPA) in the Mexican clade (Fig. S4 and Fig. 4). Using the Colombian samples with geo-referenced information, we explored the genomic divergence within the CO_Andean and CO_Caribbean subgroups. We did not identify significant (*p* > 0.05) isolation by distance (IBD) within these subgroups, as expected under rampant mobility (Fig. 5).

**Fig. 4** Maximum likelihood phylogenetic tree showing the relationship among 91 genetic pure (>80% of ancestry) avocado accessions from CGB, reference materials (RM) of Mexican (ME), Guatemalan (GU), and West Indian (WI) races, and the *P. schideana* outgroup based on 3899 SNPs. The values on nodes are the percentage of the bootstrap values for 1000 replicates. ME: Mexican race, GU: Guatemalan race, WI: West Indian race, CO: Colombian genetic group. Andean: CGB genotypes from the departments of the Andean region of Colombia, Caribbean: CGB genotypes from the departments of the Caribbean region of Colombia.

**Fig. 5** Geographic location of the Avocado Colombian genotypes from CGB with passport data. The color of the samples represents the assignation of these samples to one specific genetic cluster (CO_Andean and CO_Caribbean, West Indian, or hybrids/admixtures) based on the Phylogenetic and Ancestry analysis.

### Mixed Ancestry was Consistent with Protogynous Dichogamy

The CGB conserves avocado genotypes with ancestry to all avocado genetic races; 55.6% had a complete ancestry for the five recovered clusters in previous analyses. The majority of CGB samples have a genetic ancestry of CO_Andean (34.3%) and CO_Caribbean (13.4%), and a minor proportion of genotypes of WI (4.5%), ME (1.5%), and GU (1.5%). The other 44.4% of the genotypes that conform to this collection were admixtures. This result supported the avocado’s floral biology behavior, where the female function precedes the male function, known as protogynous dichogamy, which minimizes selfing and boosts genetic admixture. The admixture pattern was diverse; the highest proportion had a mixed ancestry of WI x G (9.7%), WI x CO_Andean (6%), and CO_Andean x WI (5.2%) (Fig. 6a and Table S4). In comparison, in the EC, most samples (75.7%) presented a pure ancestry for all genetic groups except the ME. In this collection, the genotypes with pure genetic ancestry were distributed in CO_Caribbean (37.7%), CO_Andean (11.3%), GU (23.4%), and WI (3.5%) groups; the remaining are admixtures (24.24%) where the major proportion had an ancestry of CO_Caribbean x CO_Andean (13.4%) and CO_Andean x CO_Caribbean (4.3%) (Fig. 6b and Table S4).

**Fig 6.** Barplot ancestry analysis results implemented in the CGB and EC germplasm using genotypes with complete ancestry backgrounds of Mexican (ME), Guatemalan (GU), West Indian (WI), CO_Andean, and CO_Caribbean Colombian (CO) genetic clusters as references. **(a)** CGB ancestry analysis barplot. **(b)** EC ancestry analysis barplot.

## Discussion

In this study, we analyzed the Colombian avocado genotypes coming from two collections: the AGROSAVIA germplasm bank (CGB) that preserves natives and criollos avocados and known commercial varieties (*e.g.*, ZUTANO, BACON, FUERTE, HASS), and an elite collection (EC) of avocados rootstocks from the main Colombian avocado-producing region, the Antioquia province (Table S1). Genome-reduced sequences of the Colombian germplasm (CGB) were compared with the genomic resources of avocados of Mesoamerican origin with known ancestry for the Mexican (ME), Guatemalan (GU), West Indian (WI) races, and hybrids among these races. The conducted analyses during this research classified some CGB and CWC accessions in the recognized races but the majority conformed a new Colombian genetic group, subdivided geographically in an Andean group and in a Caribbean group. The results supported that Colombia has an unexplored genetic diversity that can be used for the improvement of the crop.

### Evolutionary history and dispersion of avocado dates to the Pleistocene

Our study recovered the three primary races previously described and provided evidence to support the hybridization and introgression among races (CAÑAS-GUTIÉRREZ et al. 2019; Talavera et al. 2019). However, an unexpected result was discovering a new hotspot of avocado diversity corresponding to an exclusively Colombian genetic cluster independent of the three recognized avocado races. The Colombian genetic pool is conserved in the national germplasm bank with accessions collected from the Andes and Caribbean region. Demographic analyses supported a scenario without introgression with two parallel splits. The divergence between CO and WI predated the separation of the other two Mesoamerican races from middle-high altitudes (ME and GU). The 95% confidence intervals of the timing when all three splits occurred overlap, suggesting that the evolution of avocado, long-distance dispersal, and adaptative divergences that created the current genetic clusters occurred very fast during the climatic fluctuations of the Pleistocene, a period from 2.5 million to 11,700 years ago. At that time, the Central American isthmus had already uplifted completely, joining the North American and South American continents, helping the massive migration interchange of fauna and flora among continents, as well as human migration (Galindo-Tovar et al. 2008). Thus, our data support ecological adaptation to diverse habitats mediated by dispersion by big mammals such as the giant ground slots that inhabited the Americas during the Pleistocene. These species probably consumed avocados, dispersing seeds at long distances, generating isolation of specific populations across long periods before any human did (Diamond 1999; Barlow 2002; Chen et al. 2009). Also, our demographic model supports the dispersion of seeds and propagules by spontaneous growth from leftovers of the first hunter-gatherers’ humans that arrived in America after the last glacial maximum (LGM), ~16,000 to 13,000 years ago. They probably consumed avocados in Mesoamerica as they migrated southward (Wiersum 1997; Diamond 1999). Consequently, our demographic genetic evidence supports the hypothesis that tropical trees such as avocados may have played a unique role in the development of American cultures (Galindo-Tovar and Arzate-Fernández 2010). Avocados could be one of the first trees recurrently used *in situ* across different places in the Neotropics, promoting sedentary life and agriculture development (Smith 1966, 1969; Galindo-Tovar and Arzate-Fernández 2010).

The best demographic genetic scenario and its geographic distribution match archeological data and chroniclers’ descriptions (Galindo-Tovar et al. 2008; Galindo-Tovar and Arzate-Fernández 2010). Specifically, the preferred demographic model indicated a recent common ancestor between Mexican and Guatemalan races. Indeed, the two avocado races originated and adapted to a middle-high altitude in Mexico and Guatemala, respectively. Moreover, archeological evidence supported the *in situ* utilization of the Mexican race around 8,000 years ago in the Tehuacan Valley (Smith 1966, 1969). In parallel, our genetic data supported a recent common ancestor between the West Indian race group and the distinct Colombian cluster. Archeological and historical evidence supported that the West Indian race originated in the lowlands of Yucatan and Belize. According to Galindo-Tovar and Arzate-Fernández, (2010), migration started as follows: Maya groups living in this area utilized West Indian avocados and migrated eastward, reaching Honduras with evidence of avocado consumption by the Papayeca culture around 1,200-1,000 years ago. Also, trade activities by diverse human cultures from Mexico and Honduras extended toward Northern South America. Correspondingly, archeological evidence supports the presence of avocado seeds in the Moche Valle of Peru between 2,500-1,800 years ago and on the Peruvian Pacific coast around 1,500. Likewise, it seems that the Caral culture from Peru consumed avocados even before maize 1,200 years ago (Heiser 1979; Pozorski 1979; Skidmore 2005). The presence of cassava in Tabasco (Mexico) 4,600 years ago supported evidence for a bidirectional trading, including plant resources from the Amazon basin (Healy 1978; Pope et al. 2001; Marcus 2003).

Given this archeological evidence, northwest South America, nowadays Colombia, must have been a mandatory trading path bringing Mesoamerican and South American plant crops. The first human activities in the region date around 11,000 years ago (Oyuela-Caycedo 2008). Thus, when the Spanish arrived on the Caribbean coast of South America, they described the presence of avocado trees in Yaharo (Colombia) in 1519. Moreover, other chronicles described avocados in Peruvian Amazonia in 1542 and Ecuador in 1748, matching West Indian characteristics (Galindo-Tovar and Arzate-Fernández 2010). Morphological convergence of the avocado fruit parsimoniously supports a common ancestor between the West Indian race and the Colombian group, which opens new questions for future studies. For instance, did the pre-adaptation of the West Indian race from Central America enable the colonization of South America? After colonization, did the West Indian race radiate in South America, and did it readapt to new local conditions isolating the Colombian group? Did other races reach South America and hybridize with the West Indian race? Future research to test the above hypotheses will require extensive sampling within Colombia and surrounding areas, covering all the avocado distribution in northern South America, including Ecuador and Peru. We hypothesize that as this study supported a new genetically differentiated subpopulation in Colombia, expanding characterization to those countries could reveal other primary or secondary centers of diversity.

The novel subpopulation from northern South America indicates that previously unexplored avocado resources may have persisted via divergent selection across isolated pockets of cryptic diversity at secluded hills and valleys of the northern Andes and the Caribbean savanna. Therefore, further habitat-based population-guided collections (Castañeda-Álvarez et al. 2016) are key to prioritizing the conservation of avocado genetic resources and better characterizing these isolated pockets of diversity. Notably, we recommend new sampling trips at the foothills of the Sierra Nevada de Santa Marta, the massif of Montes de María, and the vicinity of Santa María la Antigua del Darién (southeast of the Isthmus of Panama). The former corresponds to an isolated mountain range separate from the Andean range in northern Colombia, where avocados were first described during colonial times (Reyes-Herrera et al. 2020). The latter corresponds to the first settlement founded by conquistadors in mainland America (*ca.* in 1510), initially named Dariena but abandoned a couple of decades after due to the unforgiving climate and the fierceness of the indigenous people. All three are likely sources of isolated variation for avocados, without excluding other similar hotspots of cryptic avocado diversity in Ecuador and even south of the Huancabamba depression, a critical biogeographical barrier for Andean plant taxa in the northern Andes of Peru (Weigend 2002).

### Novel genetic resources will boost avocado breeding

Modern platforms for allelic discovery (Hickey et al. 2017) would benefit by targeting cryptic pockets of genetic diversity and habitat-based population-guided collections (Castañeda-Álvarez et al. 2016), as suggested here for avocado resources in the northern Andes. This way, avocado rootstock pre-breeding efforts will not rely exclusively on exogenous diversity, likely without sufficient pre-adaptations to the local conditions of northern South America. Ultimately, the identification, conservation, and utilization of novel adaptive sources among native avocados and related wild species will enable diversifying rootstock selection by offsetting the winnowing effect (McCouch 2004) of clonal tree propagation in natural genetic variation (Ingvarsson and Dahlberg 2019).

Diversifying rootstock selection of multi-clonal genotypes and half-sib families may be boosted by genome-enabled predictions targeting discreet subpopulations such as the Colombian cluster described in this study. Multi-clonal and seedling rootstock breeding from local genetic resources could broaden the genomic basis of avocado adaptation in regions with contrasting and complex ecologies, such as the northern Andes. Families and farmers keep native avocado trees in their backyards as traditional orchards and living fences (Galindo-Tovar et al. 2008). These old avocado trees may also source novel rootstock adaptations so far unseen in current commercial plantations. Yet, satisfying the growing demand for uniform superior rootstocks is a major requirement at avocado *cv.* Hass nurseries, for which diversified clonality may be more appealing (Ingvarsson and Dahlberg 2018). Even in this case, cryptic Colombian avocado diversity may provide the desired adaptative allelic combinations to be clonally propagated in elite rootstock genotypes.

Meanwhile, rootstock breeding should be parallelized with genetic screening for innovative fruit quality traits to modernize and diversify the fresh avocado market beyond Hass. It will not be surprising that cryptic avocado gene pools and natural segregation continue sourcing new fruit varieties, as demonstrated by the unexpected discovery of Hass-like cultivars such as Carmen, Gem, Gwen, Maluma, and Méndez (Kremer-Köhne 1999). Regional consumption in Central and northern South America may even offer unexploited targets for non-Hass fruit types because consumers in those regions are historically used to marketing “criollo” avocados. For instance, market preference in northern South America tends to prioritize West Indian large and watery fruits, many of which are unknown outside local villages.

### Bridging genomic divergence with morphological differentiation

Morphological analyses in fruit tree crops like avocados are fundamental to corroborate the horticultural distinction among differentiated genetic clusters. Each avocado race exhibits specific characteristics in its morphology, phenology, ecology, and adaptation (Barrientos Priego 2010). However, race populations could be geographically isolated across different agroclimatic conditions without notorious phenotypic differentiation (Campos-Rojas et al. 2008), as expected at the initial phases of allopatric divergence (Coyne and Orr 2004). Definitory morphological characteristics appropriate for the Colombian population may offer a novel racial subdivision. López-Galé et al. (2022) suggested using fruit and seed traits to discriminate seed-donor “criollo” avocados. Oil quality could also be an indicator to discern between Colombian genotypes (Campos-Rojas et al. 2008).

Even in the absence of morphological distinctiveness, unique genetic diversity from the Colombian population may offer standing variation for adaptation to other regions where avocado plantations are expanding, as well as to cope with *in situ* climate change effects (Aitken and Whitlock 2013). Oncoming studies may rely on species distribution modeling (SDM) using current and forecasted climate data and herbaria specimens to predict the future potential distribution of avocado populations (Franklin and Miller 2010; Peterson et al. 2011). This approach will enable the prediction of the most suitable environmental conditions for avocado plantations given the future conditions, as part of an assisted gene flow platform targeting long-lived perennial crops such as fruit trees (Aitken and Bemmels 2016). Current ecological preferences (temperature, humidity, altitude) for planting avocado races (Ashworth and Clegg 2003) may be uncoupled with the most dramatic climatic scenarios. Abiotic tolerances may as well be uncoupled under changing climate, and the Colombian population could offer new tolerances so far. For instance, the Colombian group, closer to the West Indian lowland race better adapted to humid tropical areas, may offer a reservoir of resistance to *P. cinnamomi* for rootstock selection to overcome soil-borne diseases (Gross-German and Viruel 2013). In short, even though this study is the first to unveil the Colombian avocado population as a genetically distinct cluster, its phenotypic novelty is yet to be determined. Meanwhile, underutilized allelic combinations within the Colombian cluster may source avocado rootstock adaptive breeding and cultivar diversifying selection.

### Conclusions

Avocado is one of the leading exported fruit tree crops worldwide, whereas avocado *cv.* Hass is the most exported variety to the United States and other European countries. In Colombia, the native type is the most consumed internally and is considered essential for food security and a business option for communities of producing regions. In this research, we analyzed the genetic diversity of avocados conserved in the Colombian germplasm bank and cultivated materials from the primary avocado producer Colombian region (Antioquia). These resources were compared with reference varieties of the well-described Mexican, Guatemalan, and West Indian races. This study identified two ancient Colombian genetic clusters beyond the three traditionally recognized races. These Colombian subpopulations are in two distinct geographic regions, the Andes, and the Caribbean, which may reflect divergent local adaptation after the initial colonization from Mesoamerica during the Pleistocene. The exploration of the avocado genetic resources present in South America would allow identifying genotypes with superior characteristics adapted to diverse agroecological areas, which in the future may source the selection of new cultivars and rootstock genotypes.

This study is also available in Spanish for further discussion within Colombia and the region, in the same native language of researchers and producers (Supplementary File S1).

## Acknowledgments

We thank the owners and technicians of the eight avocado “Hass” orchards for donating rootstocks’ tissue, and the field assistants H.M Arias, J.M. Bedoya, K.Y. Calle, L.E. Cano, E. Carranza-Hernández, S.A. Guzmán, J.A. Henao, L.M. Mejía, A.M. Otálvaro, A.N. Sánchez and H.D. Yepes for sampling roots during 2015-2017, in collaboration with V. Velásquez-Zapata and L. Patiño. The overall Sistema General de Regalías-funded project, under which rootstocks (EC) were sampled, would have not been possible without the vision of M. Londoño, J.M. Cotes, and C.M. Gómez-Osorno, P.J Tamayo-Carmona, L.N, Martínez-Caballero, and the collaboration from G.P. Cañas-Gutiérrez, A.P. Clavijo, M. Casamitjana-Causa, O.A. Delgado-Paz, C.A. Díaz-Diez, J. Díaz-Montano, C.M. Holguín, L. Muñoz-Baena, P.E. Rodríguez-Fonseca, T.M. Rondón-Salas, and S.M. Sepúlveda-Ortega. An early version of this work was improved thanks to insights from A. Barrientos, J.I. Hormaza, M.F. Martínez, P. Manosalva, J. Patel, and G. Wilkie during the IX World Avocado Congress held on September 2019 in Medellín (Colombia). Finally, we thank the team that maintains the avocado genebank A. Caicedo, A. Hernandez, E. Rodriguez, and D. Cañar for their dedication that motivates this work.

## Author contribution

NA, CSI, YR, and RHP designed the sampling elite and CGB collection, and NA led tissue elite collection. BCJA, and DP performed DNA extraction and genomic library preparation. BCJA carried out bioinformatic analyses to discover SNPs and filtered and prepared input datasets. BCJA, CAJ, LHF, CSI, RHP, and YR made data analyses and interpreted results. All authors wrote and approved the submitted version.

## Declarations

### Data Archiving Statement

This published article and supplementary material include all data generated or analyzed during this study. The data for this study was submitted to NCBI under BioProject: PRJNA878519, which contains 384 raw Illumina sequencing data. Published sequencing data used in this study from Talavera et al. (2019) were from NCBI PRJNA564105 number. The sequences of *P. schideana* used as an outgroup were downloaded from NCBI with SRR8295605 accession number published by Rendón-Anaya et al. (2019).

## Figure legends

**Fig. 2** Pairwise F_ST_ **(a)** and bidirectional migration rates (Nem) **(b)** values among the four Avocado genetic clusters determined by population genetic structure analysis. In the Figure, GU Cluster corresponds to the Guatemalan genetic group, CO Cluster corresponds to the Colombian group, ME Cluster genetic corresponds to the Mexican genetic group, and WI Cluster corresponds to the West Indian genetic group.

## Supplementary Information

**Table S1.** List of Colombian Germplasm Bank (CGB) accessions and Elite collection (EC) genotypes *of Persea americana* Mill sequenced and analyzed in this study.

**Table S2.** OptCluster results from the analysis of the possible number of genetic clusters in the Avocado Dataset 1.

**Table S3.** Clustering assignment and ancestry proportions of the four major avocado clusters (and a fifth minor cluster) to each of the three races and the two origins (Colombian Germplasm Bank – CGB and Elite collection – EC). Clustering assignment using the optimal algoritm AGNES (Agglomerative Nesting) validated by *Optcluster* and *Ncbclust* R-packages.

**Table S4.** Summary results of the Ancestry Analysis implemented in Colombian Germplasm Bank (CGB) and Elite collection (EC) avocado genotypes.

**Figure S1.** The eight demographic evolutionary scenarios used in DIYABC random forest to test the scenario that best fit with the SNP polymorphism among the three recognized avocado races, Mexican (ME), Guatemalan (GU), and West-Indian (WI), and the Colombian group (CO) found in this study. The time in the scenarios is measured in generations and is not to scale. **(a)**. Scenario 1 was the null hypothesis, indicating that all the four genetic groups diverged simultaneously at *t*_1_. Scenarios 2-4 considered three divergence times among the four groups (i.e., *t*_1_, *t*_2_, and *t*_3_) but tested what was the most ancient group, considering the two extremes of the distribution, from Mesoamerica to the North of South America. In scenario 2 **(b)**, the most ancient divergence occurred at *t*_3_ and derived CO. Then, at *t*_2_ derived WI, and finally, at *t*_1_, the most recent common ancestor between ME and GU diverged. In scenario 3 **(c)**, again, CO is derived from the most ancient divergence *t*_3_. Then, at time *t*_2_ evolved GU, and at the most recent divergence time, *t*_1_ evolved ME and WI from the most recent common ancestor. Finally, in scenario 4 **(d)**, the ME group is the most ancient group derived at *t*_3_. Then, at time *t*_2_ derived GU; and at *t*_1_ diverged CO and WI from the most recent common ancestor. Scenario 5 **(e)** contemplated a potential hybrid evolution. Here, CO is the most ancient group derived at *t*_3_; then, at *t*_2_, WI and ME split; finally, at *t*_1_, GU evolved from the hybridization between WI and ME. Alternatively, scenarios 6-8 simulate the evolution of the most recent common ancestor at *t*_3_ from the four groups, but then the parallel evolution of two clades that split at times *t*_2_ and *t*_1_ generating two sister groups. In scenario 6 **(f)**, the sister groups were WI-ME, which split at *t*_2_, and CO-GU that split at *t*_1_. In scenario 7 **(g)**, the sister groups were GU-ME, which split at *t*_2_, and CO-WI, that split at *t*_1_. Finally, in scenario 8 **(h)**, the sister groups were WI-GU, which split at *t*_2_, and CO-ME, that split at *t*_1_.

**Figure S2.** Metrics determined in the Avocado SNPs detected in dataset 3. **(a)** Mean Depth of Sequencing - MDS, **(b)** Minor Allele Frequency - MAF, **(c)** Polymorphic Information Content - PIC, **(d)** Genetic Diversity - GD, **(e)** Nucleotide Diversity - ND and **(f)** Observed Heterozygosity - Ho.

**Figure S3.** Mantel correlations between genetic differentiation and geographic distance within and between pairs of clusters (GU Cluster, CO Cluster, ME Cluster, and WI Cluster). Diagrams depict pairwise F_ST_/(1–F_ST_) vs. geographic distance (Km). Each subpanel highlights explicit comparisons of each cluster against the Guatemalan - GU **(a)**, Colombian - CO **(b)**, Mexican - ME **(c),** and West Indies - WI **(d)** genetic clusters, according to Fig 2. Therefore, clouds of dots at the bottom of each subpanel correspond to within cluster comparisons. Dots are colored following Fig. 2 and Table S3. Trend lines are not displayed because Mantel tests were not significant (*p-value* > 0.05).

**Figure S4.** Maximum likelihood phylogenetic tree showing the relationship among 203 avocado accessions from CGB, reference materials (RM) of Mexican (ME), Guatemalan (GU), and West Indian (WI) races, and the *P. schideana* outgroup based on 3899 SNPs. The values on nodes are the percentage of the bootstrap values for 1000 replicates. ME: Mexican race, GU: Guatemalan race, WI: West Indian race, CO: Colombian genetic group.

**Supplementary File S1.** The manuscript in Spanish.

